# SAR324 and related lineages are associated with the evolutionary history and origins of dsr-mediated sulfur oxidation

**DOI:** 10.1101/2024.01.30.576480

**Authors:** Katherine M. Klier, Cody Martin, Marguerite V. Langwig, Karthik Anantharaman

## Abstract

Microorganisms play vital roles in sulfur cycling through the oxidation of elemental sulfur and reduction of sulfite. These metabolisms are catalyzed by dissimilatory sulfite reductases (dsr) which function in the reductive (dsr) or reverse, oxidative direction (rdsr). Dsr-based sulfite reduction is an ancient metabolism and has been proposed to fuel energy metabolism in some of Earth’s earliest microorganisms. Conversely, sulfur oxidation is believed to have evolved later in association with the widespread availability of oxygen on Earth. Organisms are generally believed to carry out either the reductive *or* oxidative pathway, yet a subset of bacterial phyla have recently been discovered with gene combinations that implicate them in both pathways. A comprehensive global investigation into the metabolisms of these phyla regarding dsr can shed light on the evolutionary underpinnings of sulfur metabolism but is currently lacking. In this study, we selected one of these phyla, the abundant and metabolically versatile candidate phylum SAR324, to study the ecology and evolution of dsr and rdsr. We confirmed that phylogenetically, environmentally, and geographically diverse SAR324 contained dsr, rdsr, or both. Comprehensive phylogenetic analyses with other dsr-encoding bacterial and archaeal phyla revealed that organisms encoding both dsr and rdsr genes are constrained to a few phyla, which we term “transitionary clades for sulfur oxidation”, and these phyla are phylogenetically positioned at the interface between well-defined oxidative and reductive bacterial clades. Together, this research suggests that SAR324 and other transitionary clades are associated with the evolutionary history and origins of the reverse dsr pathway in bacteria.

## INTRODUCTION

Sulfur is ubiquitous in the Earth’s atmosphere, lithosphere, and hydrosphere (1–3). Sulfur is essential to life and comprises key components of amino acids such as cysteine and methionine, organic compounds, and cofactors (4–6). The sulfur cycle constitutes a major biogeochemical cycle on our planet and actively interacts with other biogeochemical cycles such as those of carbon, nitrogen, and iron (7–9). For example, transformations of sulfur compounds such as sulfate and sulfide are closely associated with greenhouse gases such as methane and carbon dioxide, which are vital to monitor in the face of climate change (10). Microorganisms play a critical role in the sulfur cycle, primarily by using sulfur compounds as electron donors or acceptors for energy metabolism (11–14). Nevertheless, the accumulation of some reduced sulfur compounds, such as hydrogen sulfide, can have toxic effects on environments (15,16), highlighting the importance of microbial sulfur metabolism.

Microbial sulfur metabolism is diverse and consists of several metabolic pathways, including sulfur assimilation, dissimilation, and disproportion (17). A metabolism of particular interest that has garnered significant attention from the scientific community is dissimilatory sulfite reduction (dsr) (18–21). Predicted to have evolved over 3 billion years ago, dsr metabolism is thought to have been present in some of earth’s most ancient microbial lifeforms (20,21). In recent years, the number of microorganisms known to possess genes for dsr metabolism has grown substantially (18,19).

In the forward direction, dsr catalyzes sulfite reduction, a crucial intermediate step in the reduction of sulfate to sulfide, providing energy to organisms in anoxic environments (22,23). The reverse-dissimilarity sulfite reductase pathway, often predicted to have evolved after the forward pathway, can oxidize elemental sulfur to sulfite (21,24–26) and is a crucial part of sulfur oxidation metabolism. Therefore, it has also been termed the oxidative dsr pathway.

Numerous dsr proteins exist, some of which have been associated with the direction of the sulfur transformation reaction that an organism catalyzes (i.e. reductive or oxidative dsr pathway). Key dsr enzymes involved in both pathways include dsrAB, the catalytic core (27,28), and dsrC which acts as a physiological partner and “redox hub” in dsr activity (29). DsrD, recently identified as an allosteric activator of dsrAB sulfite reductase activity has been used as a marker for the reductive pathway (18,30). DsrEFH are thought to act as a sulfur carrier to dsrC and, thus, are typically associated with microbial sulfur oxidation (18,31). DsrL has been shown to be essential for sulfur oxidation but is also present in sulfite reducers (32). Additional proteins that may be present include dsrM, dsrK, dsrJ, dsrO, dsrP, and dsrR, which perform various functions, ranging from posttranscriptional control to the electron transport chain (33,34). Dsr proteins are encoded for by genes of the same name.

Phylogeny, along with gene content, has also been used to assess pathway directionality, with three main divisions, archaeal reductive, bacterial reductive, and bacterial oxidative typically being described (19,35). Likewise, microorganisms in general that metabolize sulfur are often described as *either* Sulfur Reducing Microorganisms (SRMs) or Sulfur Oxidizing Microorganisms (SOMs) (9,36). Common examples of SRM’s are from the phyla Desulfobacteria, Bacillota (formally Firmicutes), Nitrospirota, and Thermoproteota (formally Crenarcheaota) (37,38). Examples of SOMs are organisms from the phyla Chlorobi (now classified under the phyla Bacteroidota) and Pseudomonadota (a phyla previously classified as proteobacteria encompassing Gamma, Alpha, Beta, and Muproteobacteria) (39–41).

Recent research has challenged traditional classifications, revealing certain phyla with both reductive and oxidative *dsr* genes in the same genome. For example, the coexistence of genes encoding for dsrD and dsrEFH in genomes belonging to phyla such as SAR324 and Actinomycetota has made it difficult to discern whether these organisms participate in the reductive or oxidative pathway. It has been suggested these organisms have the potential to switch between reductive and oxidative sulfur cycling pathways. On the other hand, the incomplete nature of many genomes, recovered as Metagenome Assembled Genomes (MAGs) and Single Cell Amplified Genomes (SAGs), has also raised concerns about mis-binning or mis-assembly as an explanation for unique gene combinations (18,19). With only a handful of these genomes being described in literature, it has remained unclear if these unique gene combinations are a biological phenomenon (as opposed to mis-assembly) and, if so, why might these phyla carry them.

Inspired by these discoveries, we delved further into our exploration, zeroing in on an intriguing phylum: Candidate Phyla SAR324, hereafter referred to as SAR324. Thriving in diverse marine environments, SAR324’s metabolically versatile nature adds complexity to its role in sulfur cycling (42–46). Not only do select SAR324 genomes exhibit unique *dsr* gene combinations, but their dsrA sequences have been found to stand apart phylogenetically from other bacterial sulfur oxidizers and reducers (45). These distinctive features make SAR324 a compelling subject for unraveling the intricacies of dsr-mediated sulfur transformation.

In this study, we undertake a comprehensive analysis of dsr in SAR324, leveraging all publicly available genomes. We describe both dsr and oxidative dsr pathways and shed light on the unique phylogeny of SAR324 dsrAB sequences positioned between oxidative and reductive types. Our findings constrain the evolution of the oxidative dsr pathway and sulfur oxidation metabolism to SAR324 and closely related lineages and reveal an undescribed combination of reductive and oxidative genes in the genomes of these same lineages.

## METHODS

### Data acquisition

To create a SAR324 genome database, the key word “SAR324” was searched for in the National Center for Biotechnology Information (NCBI) database (47) and Genome Taxonomy Database (GTDB) (48). Accession number hits were used to obtain and download genomes from NCBI GenBank. Additional genomes were obtained from the Joint Genome Institute (JGI) (49) database. Genomes from the JGI database had been previously published (43). Genomes from phyla Aquificota were downloaded from NCBI GenBank and included in this dataset to root the phylogenetic tree. SAR324 metadata can be found in **Supplemental Table 1**.

To create a comprehensive dataset of microbial and archaeal sulfur cycling microorganisms for which to search for dsr homologues, 4631 additional publicly available genomes were obtained from the NCBI GenBank database. Microbial and archaeal phyla associated with dsr-sulfur cycling were chosen based on previously published literature (18,50,51). Our SAR324/Aquificota dataset was also included in this dataset. In total, the original database consisted of 4856 genomes. Genome identifiers for the dataset are located **Supplemental Table 2**. This table exclusively lists genomes that were included in the downstream analyses.

### Genome classification

Taxonomic classification/verification of genomes was performed using GTDBtk v2.1.1 (52) with the 214-release database using the “--wf” option.

Genome identifiers and their GTDBtk classifications can be found in **Supplemental Table 2**.

### Genome dereplication and quality checking

Genome dereplication for the comprehensive database of microbial and archaeal sulfur cycling microorganisms was completed using dRep v3.3.0 (53) with an average nucleotide identity of 95% to produce optimal species-level representative genomes (54). During dereplication by dRep, genomes were filtered for medium-to high-quality draft genomes using checkM v1.2.0 (55) with a 50% completeness and 10% contamination threshold per established minimum standards (56). dRep was run with default parameters and the following flags: “-nc 0.5 -con 10 -comp 50”.

Non-dereplicated genomes from the SAR324 analyses were quality checked with checkM v1.2.0 with the “lineage_wf” workflow and default parameters. Genomes that did not pass the completeness and contamination thresholds were removed from analyses.

Completeness and contamination estimates for both datasets can be found in **Supplemental Table 3**.

### Identification of dsr and ribosomal subunit protein homologs

Generation of HMM datasets used in this paper are described elsewhere (18,42,57). To identify ribosomal protein subunit homologs, the open reading frames (ORFs) for each genome were predicted using Prodigal v2.6.3 (58) and used to identify 16 ribosomal subunit protein sequences (L2, L3, L4, L5, L6, L14, L15, L16, L18, L22, L24, S3, S8, S10, S17, S19) with HMMER v3.3.2 (59). HMMER was used with default parameters and the following flag: “-E 1e-5”. To search the genomes for dsr homologs, ORFs were predicted using Prodigal v2.6.3 with default parameters, and ORFs were searched against a custom database of dsr proteins (dsrABCDEFHMKJOPLSRT) using HMMER v3.3.2 with the “--cut_tc” flag.

### Maximum likelihood ribosomal protein phylogeny

The SAR324 phylogeny (**Figure 1B**) was constructed with 162 medium-to-high-quality SAR324 genomes. Ribosomal protein subunits were identified as described above. Individual alignments for each ribosomal subunit protein were produced using MAFFT v7.490 (60,61) with default parameters and then uploaded to Geneious v2022.0.2 (62) for curation. Duplicate hits to the same ribosomal protein unit within the same genome were removed by manual inspection. The individual alignments were concatenated, and alignments with less than 60% of un-gapped residues from the consensus sequence were removed from the analysis. The alignment was then masked at a threshold of 97%.

**Figure 1.**
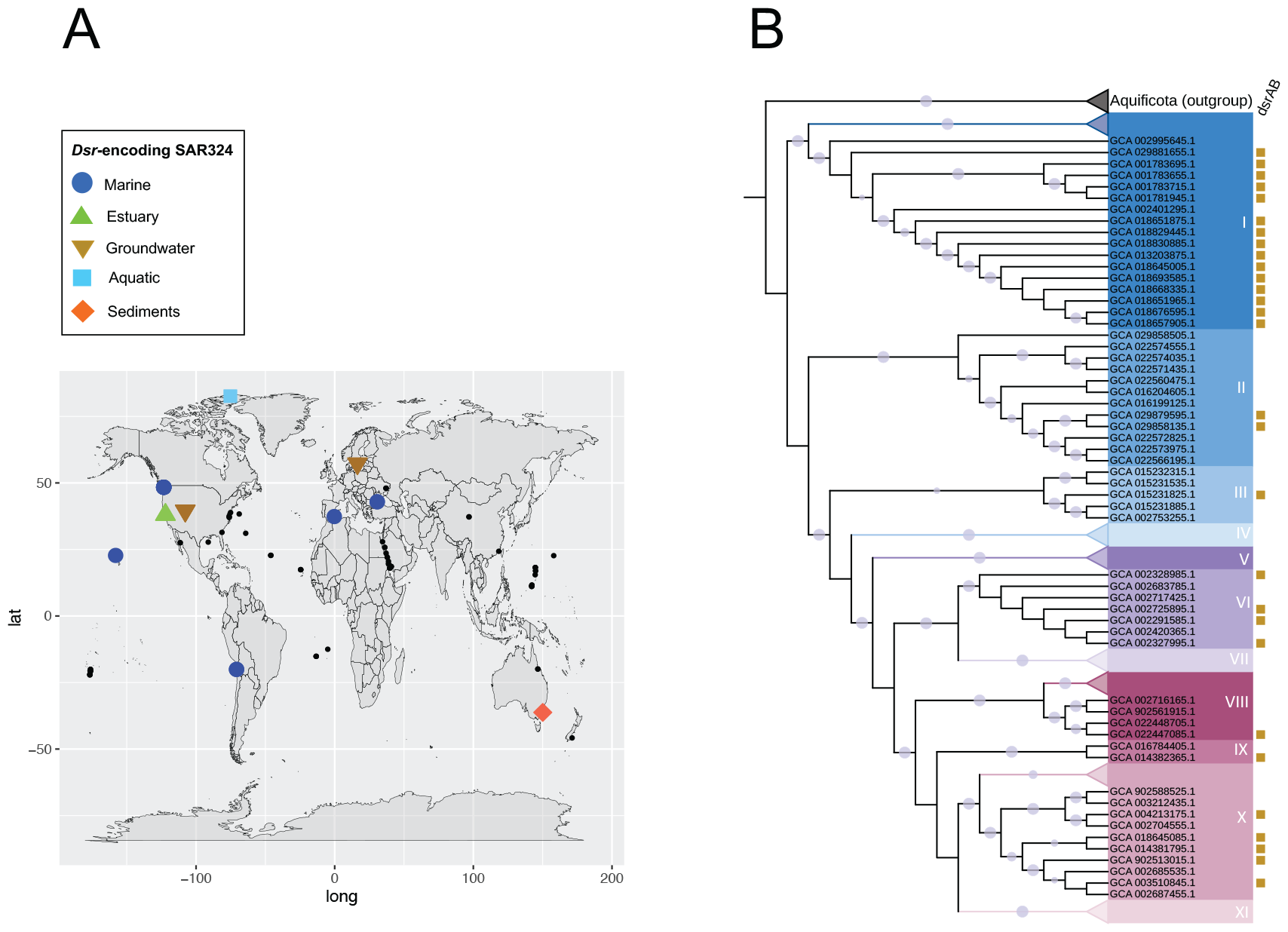
Dsr is present in globally distributed SAR324. A) Geographic distribution of dsr-encoding SAR324 genomes correlated to the ecosystem type they were retrieved from. Non-dsr encoding SAR324 genomes are included on the map and marked by small black circles. B) Maximum likelihood phylogeny of 145 SAR324 genomes generated via concatenation of 16 ribosomal protein subunits. The background colors of text in the phylogeny denote SAR324 clades I-XI. Bootstrap support >90 is annotated with a lavender circle on the branches of the tree. Yellow squares denote presence or absence of dsrAB in SAR324 genomes. Clades/subclades in which dsrAB was not found were collapsed. The tree is rooted to genomes belonging to the phyla Aquificota.

The phylogeny was created using IQ-TREE v2.1.4 (63) with the following parameters: “-T AUTO-ntmax 15 -bnni -bb 1000 -m TESTMERGE”.

The newick tree file and curated alignment are located in **Supplemental Files 1 and 2**.

### Maximum likelihood dsr gene trees

The DsrAB phylogeny (**Figure 3**) was generated by dereplicating and quality checking the 4,856 genomes downloaded from NCBI and JGI as described above. Genomes were searched for dsr homologs as described above.

Alignments of dsrAB proteins were constructed using MAFFT v7.490 with the same parameters as the ribosomal subunit phylogeny. After concatenation, only genomes that contained at least partial dsrA *and* dsrB sequences were retained in the concatenated alignment. The alignment was then masked at a threshold of 97%. For the dsrEFH tree (**Figure 4**), the same methods were utilized as the dsrAB phylogeny, with the exception that, after concatenation, only alignments with dsrE and at least one of dsrF and dsrH were kept. The same methods as the dsrAB phylogeny were also used to construct the dsrR tree (**Figure 6**), except all sequences were kept.

The same methods were followed for the SAR324 dsrAB phylogeny (**Figure 5A**) and SAR324 dsrEFH phylogeny (**Figure 5B**) as for **Figure 3** and **Figure 4** respectively, with the exception that in lieu of the larger sulfur cycling database, the SAR324 database was searched for dsr homologs.

All phylogenies were constructed with IQ-Tree v2.1.4 with the same parameters as the ribosomal subunit phylogeny. Newick tree files and curated alignments are located in **Supplemental Files 3 - 12**.

### Phylogenetic tree visualization

All phylogenetic trees were visualized and annotated using iTol: Interactive Tree of Life version 6 (64).

### Gene organization analysis

To examine the gene content and organization in dsr-encoding genomes (**Figure 2**), a custom python script was used to search ORFs against our dsr database using HMMER v3.3.2, orient and order genes based on genome position and produce preliminary gene structures (script available at https://github.com/cody-mar10/operon_finder). Simplified operon visualizations were manually generated. All genes were oriented with respect to the direction of dsrAB, including sequences found on different scaffolds. Duplicated dsr homologs in the same genome, aside from dsrAB, were omitted from **Figure 2** unless in the vicinity of a dsrAB copy. Full gene content information can be found in **Supplemental File 13**.

**Figure 2.**
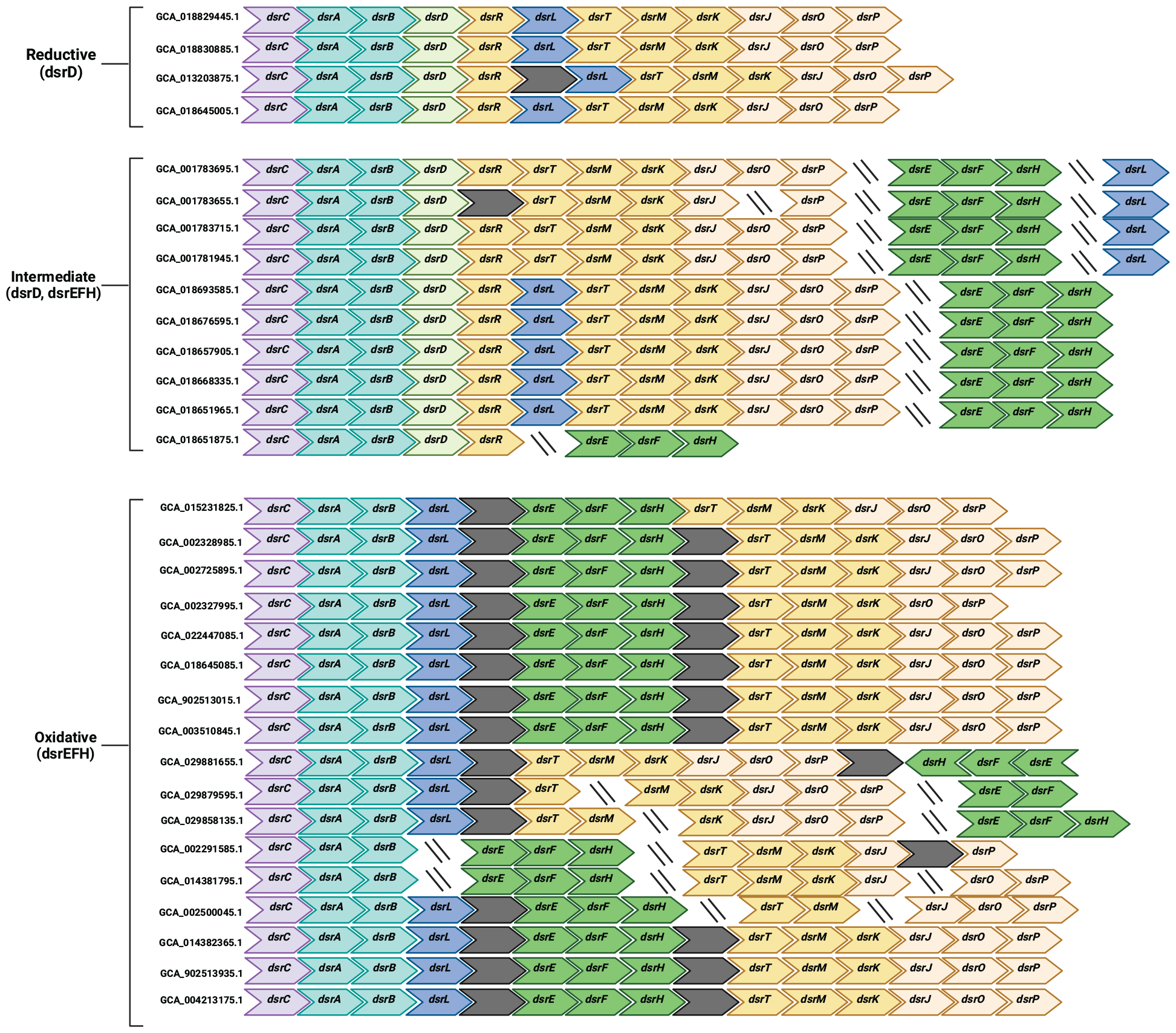
Analysis of SAR324 dsr content reveals distinct groupings. Visualization of *dsr* gene structure in 31 SAR324 genomes. Slanted lines indicate the end of a scaffold, whereas grey boxes indicate the presence of one or more hypothetical/non-dsr proteins. SAR324 genome identifiers are included to the left of each structure. Figure generated with BioRender.com

### Geographic visualization

Latitude and longitudinal coordinates for SAR324 genomes were obtained from NCBI metadata. Genomes without these metadata were omitted from **Figure 1A. Figure 1A** was generated in R v4.1.1 using the Tidyverse (65) and ggplot2 packages (66).

## RESULTS

### Geographically and phylogenetically diverse members of SAR324 encode dsr

Previous research has suggested that only a subset of SAR324 genomes contain pathways for sulfur metabolism including dsr. To explore the full diversity of dsr in SAR324, we first focused on investigating the prevalence and phylogenetic distribution of *dsr* genes in the phylum. To this end, we searched 162 medium-to-high quality SAR324 genomes for homologs to dsrA and dsrB, which constitute the catalytic core of dsr activity. Of these, 31 SAR324 genomes were identified that encode dsrAB (**Supplemental Table 1**).

To assess the geographic distribution of dsr-encoding SAR324, we plotted the location of origin for dsr-encoding SAR324 genomes for which latitude and longitude metadata was available (**Figure 1A**). We found that dsr-encoding SAR324 were globally distributed. Environments from which these genomes originated also varied and included anoxic groundwater, marine ecosystems, estuaries, hypoxic lakes, and sediment (**Figure 1A, Supplemental Table 1**).

To determine how dsr was phylogenetically distributed in SAR324, we generated a maximum likelihood concatenated ribosomal protein subunit phylogeny. The phylogeny consisted of representatives from 145 medium-to-high quality SAR324 genomes that contained sufficient ribosomal protein data. This is, to our knowledge, the most comprehensive global phylogeny of SAR324 available. The phylogeny showed that SAR324 genomes split into distinct clades, which we refer to as clades I-XI (**Figure 1B**).

Dsr was observed in phylogenetically diverse SAR324, present in clades I, II, III, VI, VIII, IX and X. We observed patterns of both vertical and horizontal gene transfer (VGT, HGT) for dsrAB among SAR324 organisms. For example, evidence of VGT was observed in a clade I sub lineage. Conversely, dsrAB were also found interspersed throughout nearly all SAR324 clades, suggesting HGT is involved. This is consistent with previous studies that have shown *dsr* genes to move via HGT and VGT (18,19,67,68).

Overall, these data suggest that SAR324 contributes to sulfur cycling via the dsr pathway in a variety of niches across the globe. Further, dsr*-*encoding SAR324 are phylogenetically diverse, suggesting that sulfur cycling in SAR324 is more widespread than has previously been suggested (44).

### SAR324 genomes encode genes for reductive type dsr, oxidative type dsr, or both forms

The data presented in the previous section confirm that dsr is prevalent in SAR324, yet, due to prior studies characterizing unique gene combinations in SAR324, the specific dsr sulfur cycling pathway (reductive vs oxidative) in which this phylum participates remained uncertain. Some previous investigations have posited SAR324’s association with the oxidative pathway (45,69). However, the limited sampling of SAR324 genomes encoding dsr in these studies introduced a challenge in establishing definitive conclusions regarding this information. To resolve these discrepancies, we analyzed the presence/absence and organization of all *dsr* genes within our dsrAB-encoding SAR324 genomes, focusing specifically on the presence/absence of *dsrD* and *dsrEFH* genes. This significantly expanded the number of dsr-encoding SAR324 genomes that have been investigated in previous studies (18,19,44,69).

The 31 SAR324 genomes encoded different combinations of dsr proteins (**Figure 2**). Notably, 10 genomes encoded both dsrD and dsrEFH, which are markers for reductive and oxidative dsr, respectively. Aside from these genomes, four SAR324 encoded dsrD but did not encode dsrEFH, suggesting they likely participate in the reductive pathway. Conversely, 17 genomes encoded dsrEFH but did not encode dsrD, suggesting these organisms participated in the oxidative pathway. Consequently, the SAR324 phylum encompasses organisms that have gene signatures of both the oxidative and reductive dsr pathway, both across and within singular genomes. These genomes were obtained from 13 different NCBI BioProjects and 5 different environment types (**Supplemental Table 1**). Hence, we posit that these observations of unique *dsr* gene combinations do indeed reflect a real biological phenomenon.

Interestingly, in the non-dsrD-encoding group, dsrEFH was typically found encoded for on the same scaffold and encoded in the same direction as dsrAB, separated only by dsrL and one or two other proteins (**Figure 2, Supplemental File 13**). In the dsrD-encoding group, dsrEFH was encoded for in a different genomic location (i.e. on a different scaffold than dsrAB). This may have implications for the reconstruction of MAGs and SAGs from the dsrD-encoding group, where researchers may have been less likely to recover both dsrAB and dsrEFH during assembly. Therefore, there is a possibility the other SAR324 that encode dsrD also encode dsrEFH but were not identified due to their separation in SAR324 genomes.

Collectively, our data suggests that SAR324 organisms can be categorized into three groups: reductive (dsrD-encoding) oxidative (non-dsrD-encoding), and an intermediate category (organisms which encode both dsrD and dsrEFH). Given that dsrEFH has been shown to work in conjunction with dsrC (31), the separation of *dsrEFH* genes in relation to those for *dsrABC* in the intermediate group may imply that these genomes lean towards a reductive metabolism, however it cannot be ruled out that they cannot switch between reductive and oxidative. These analyses further emphasize the unique metabolic flexibility and breadth of SAR324 and demonstrate. previous observations about the distinctive gene combinations in SAR324 extend to the entire phylum.

### Dsr sequences from SAR324 and several closely related phyla form a “transitionary clade” between reductive and oxidative types

To confirm if a genome is associated with oxidative or reductive type dsr, the phylogenetic placement among other known dsr-encoding bacterial/archaeal genomes is often used in conjunction with gene content analysis. Therefore, we investigated the placement of SAR324 among other dsr-encoding phyla to further determine their role in the reductive and/or oxidative pathway. We were particularly interested to know where the genomes encoding both dsrD and dsrEFH were positioned. Our approach involved construction of a concatenated dsrAB phylogenetic tree consisting of genomes from 29 bacterial and archaeal phyla (**Figure 3A**). The phylogeny contained 763 unique dsrAB sequences from 725 genomes. To help differentiate between reductive and oxidative types, we annotated genomes encoding dsrD as well as genomes encoding dsrE, which was used as a marker for the dsrEFH complex.

**Figure 3.**
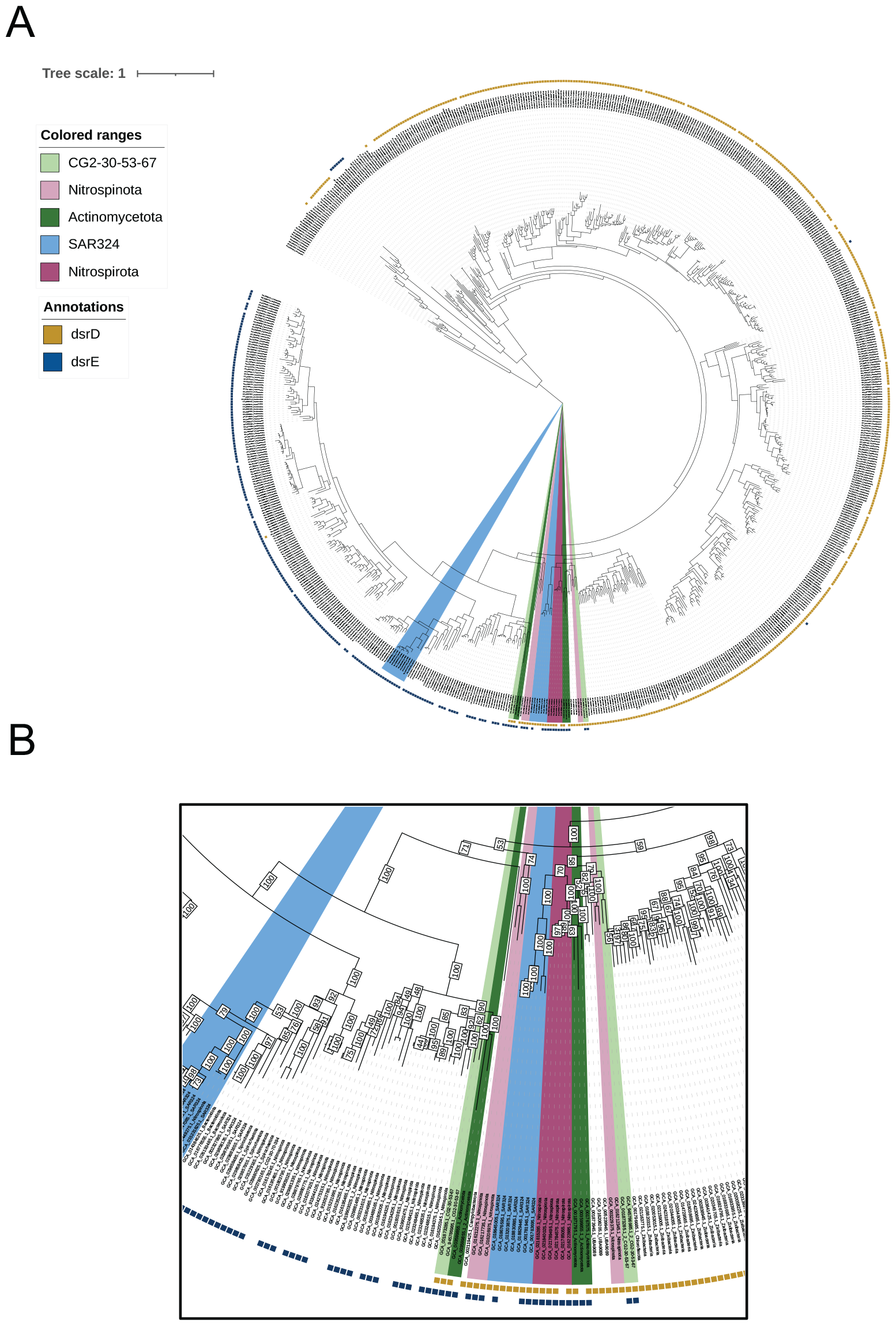
DsrAB tree reveals select genomes that bridge the reductive and oxidative types. A) Maximum likelihood dsrAB phylogenetic tree. 763 total dsrAB sequences from 29 phyla are shown. Yellow squares annotate genomes encoding dsrD while blue squares annotate those encoding dsrE. The transitionary clades are color coded by phyla. Each branch is labeled with the genome identifier and phyla which the genome belongs to. B) Zoomed in portion of the tree containing the transitionary clades and illustrating bootstraps on the tree.

As expected based on previous findings that microorganisms using reductive and oxidative dsr pathways are phylogenetically distinct (18,24,35), genomes that encoded dsrD clustered together, whereas those encoding dsrE clustered together (**Figure 3**). The exception to this was a clade of Methylomirabilota (previously Rokubacteria) which positioned itself towards the archaeal root of the tree and encoded dsrE. Among the phyla positioned with the dsrD-encoding genomes included known sulfate/sulfite reducers such as Desulfobacteria, Bacillota and Thermoproteota. Organisms that clustered with genomes encoding dsrE included known sulfur oxidizers such as Bacteroidota (previously Chlorobi) and Pseudomonadota. Additionally, the dsrD-lacking SAR324 group, which contained only oxidative type gene signatures, clustered within this latter group (**Figure 3A**).

Notably, the dsrD-encoding SAR324 group, which includes genomes encoding both dsrD and dsrE, exhibited a unique positioning. Instead of being firmly entrenched in either category (reductive or oxidative), SAR324 clustered between the reductive and oxidative bacterial types. Intriguingly, this pattern held true for other phyla that encoded both dsrD and dsrE within their genomes such as members of Nitrospinota, Nitrospirota, Actinomycetota, and CG2-30-53-67. We refer to these clades as “transitionary clades” that bridge oxidative and reductive types (**Figure 3**).

Given these findings, we examined the dsr structure of select genomes from these phyla to determine if they had patterns similar to SAR324 **(Supplemental Figure 1**). Interestingly, members of Nitrospinota and Nitrospirota contained similarities to SAR324 as, in some genomes, dsrEFH was sometimes found encoded in a different genomic location than dsrAB. Additionally, genomes from these phyla contained representatives with gene signatures across all three categories (reductive, oxidative, and intermediate) described in **Figure 2**. However, in contrast to SAR324 genomes, dsrABC, dsrD, and dsrEFH were sometimes all found encoded for on the same scaffold in Nitrospinota and Nitrospirota genomes.

Actinomycetota and CG2-30-53-67, were notable since some genomes were differentiated in that the genomes encoded two or more copies of dsrAB. In these genomes, one copy of dsrAB was phylogenetically positioned closer to sulfur oxidizers such as Bacteriodota, whereas the other dsrAB copy was closer to reducers such as Desulfobacteria (**Figure 3A, Supplemental Figure 1**). The bootstrap supports associated with the nodes at which genomes from the transitionary clades branched off from each other and other sulfur oxidizers/reducers were relatively low (<80), indicating that the position of the phyla was not well resolved as either belonging within the oxidative or reductive type clades, further solidifying them as a transition group (**Figure 3B**).

Collectively, our data suggests that members of SAR324, Nitrospinota, Nitrospirota, Actinomycetota, and CG2-30-53-67, which we collectively refer to as “transitionary clades”, exhibit unique *dsr* gene combinations within the same genome, cluster in a unique phylogenetic position, and some have representatives spanning the reductive and oxidative types. Rather than, or in addition to, being able to switch between the two pathways, we hypothesized that these phyla may have served as evolutionary intermediates during the transition from reductive to oxidative dsr bacterial types.

### Evolution of dsrEFH in the transitionary clades

Considering our hypothesis of members of the transitionary clades being involved in the evolutionary transition from the reductive to the oxidative dsr type, we could expect that they would be among the first to have evolved *dsrEFH* genes. To test this, we generated a phylogenetic tree of all dsrEFH-encoding genomes from our original dereplicated and quality filtered sulfur cycling dataset (**Figure 4**). The tree consisted of 253 dsrEFH sequences from 246 unique genomes.

**Figure 4.**
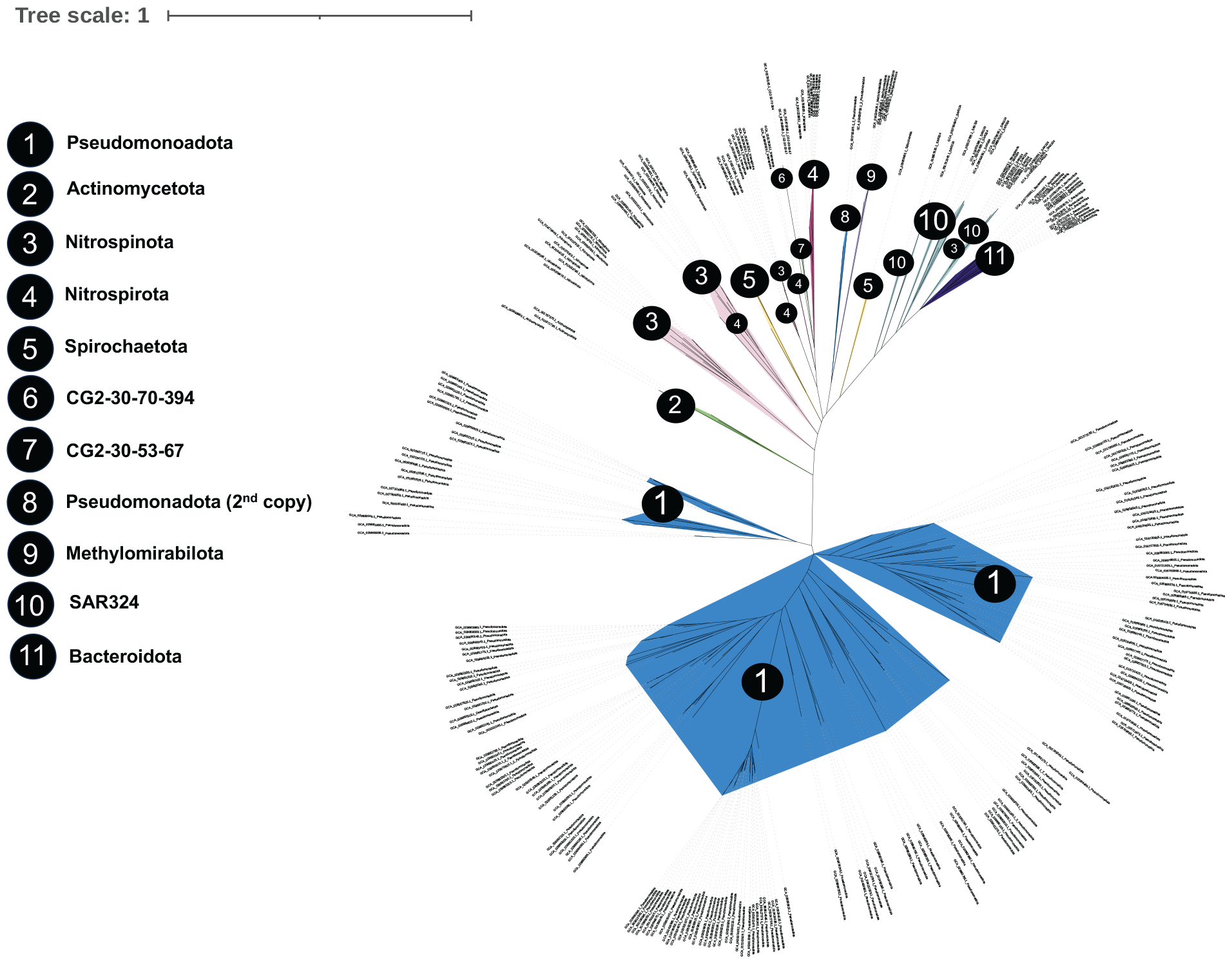
The transitionary phyla are basal in a dsrEFH gene tree. An unrooted maximum likelihood dsrEFH gene tree is shown. Genomes, including those that do not encode dsrAB are included in the unrooted tree. Genomes are color coded by phyla and numbers placed on clades are correlated to phyla and annotated in the legend. Each branch is labeled with the genome identifier and phyla which the genome belongs to.

The unrooted dsrEFH tree identified two main clades, one composed of Pseudomonadota, and the other composed of the transitionary phyla along with Bacteroidota and Spirochaetota. This observation was consistent with a previously reported hypothesis that stated oxidative dsr metabolism may have evolved in at least two separate events (26). In the latter grouping containing Bacteroidota, members of the transitionary phyla were basal with Actinomycetota and Nitrospinota being the most basal.

In summary, our data indicated that members of the transitionary clades comprising of SAR324 and other phyla, were key to the switch from reductive to oxidative metabolism, contributing to the evolution of well-known oxidizers like Bacteroidota. In contrast, the phyla most closely related to the evolution of oxidative dsr in Pseudomonadota remain unclear, given their dsrEFH’s different evolutionary history and absence of representative members that have any reductive signatures. Further sequencing of less-studied phyla may be necessary to clarify this aspect.

### Acquisition of oxidative *dsr* genes consisted of a combination of vertical and horizontal gene transfer

The data presented so far suggest that the transitionary clades played a role in the shift towards modern sulfur oxidizers. To determine if *dsrEFH* genes co-evolved with *dsrAB* genes, or if they were inherited independently, perhaps through horizontal gene transfer, we compared the dsrAB and dsrEFH phylogenies of SAR324. The paired dsrAB and dsrEFH trees were rooted to the same genomes for the clearest comparisons (**Figure 5**).

**Figure 5.**
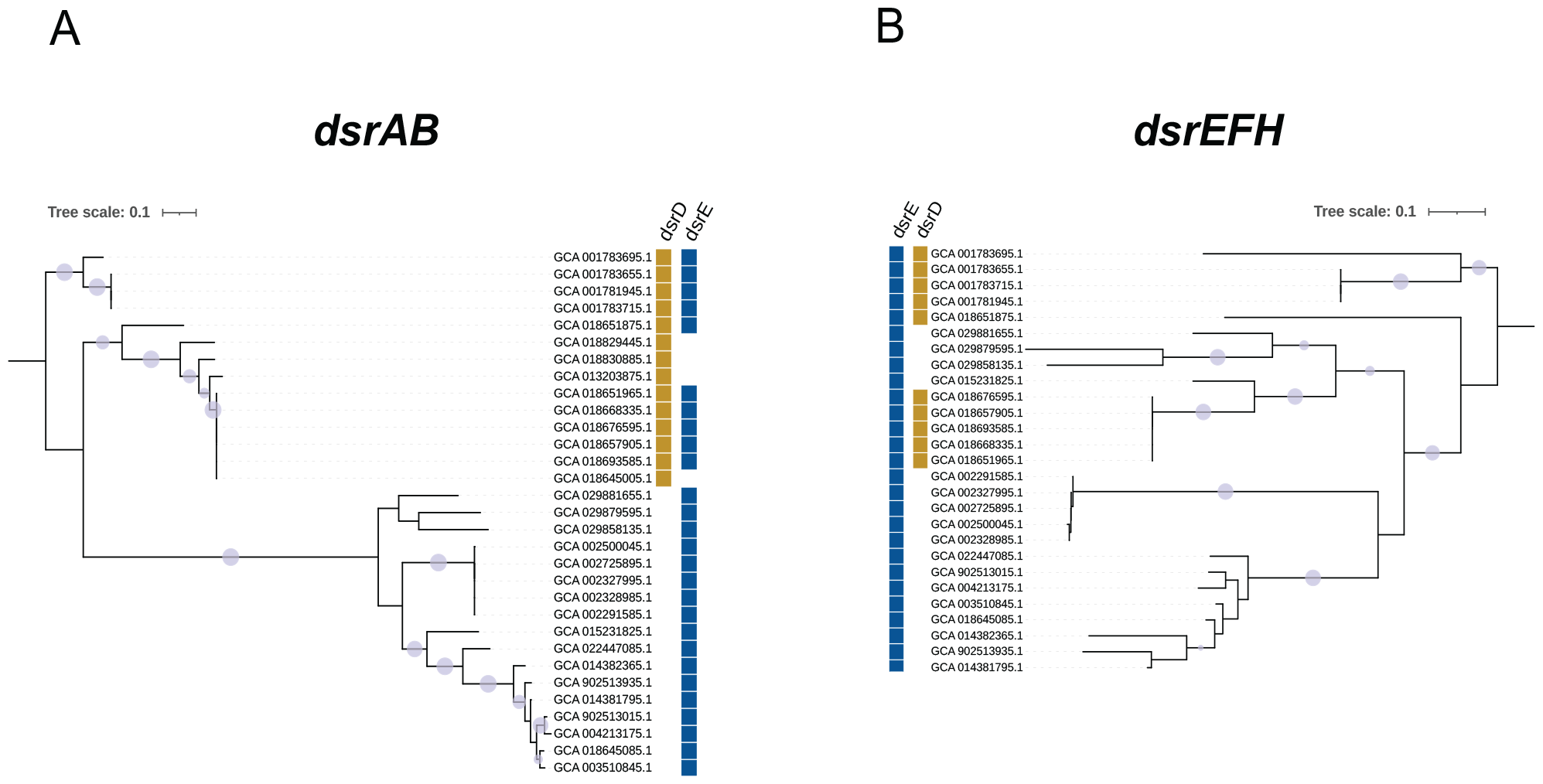
Comparison of SAR324 dsrAB (A) and dsrEFH (B) phylogenies suggests horizontal gene transfer. Maximum likelihood phylogenies are shown. Phylogenies are rooted to the same genomes. Bootstrap support >90 is shown with a lavender circle.

In some cases, there was congruent evolutionary history between the dsrEFH and dsrAB phylogenies. For example, there was some separation of the dsrD-encoding and non-dsrD-encoding genomes in both phylogenies. In addition, in both trees, some groups of genomes such as GCA_001783695.1, GCA_001783655.1, GCA_001783715.1, and GCA_001781945.1 grouped closely together in a clade. In other cases, there were discrepancies. For example, in the dsrAB tree, there was a complete separation of dsrD encoding and non-dsrD encoding genomes. However, in the dsrEFH tree, the clusters of dsrD lacking trees were permeated by a group of non-dsrD encoding genomes. Additionally, in the dsrAB tree, the dsrD-lacking clade was clearly separated by a much longer branch length, whereas this was not the case in the dsrEFH tree, and the genomes were more evenly dispersed. Therefore, like the *dsrAB* genes, it is likely that *dsrEFH* genes also move between bacteria through a combination of VGT and HGT.

If indeed HGT of *dsrEFH* has occurred, it could also explain the deep branching position of the Methylomirabilota in the dsrAB phylogeny, even though they encode dsrEFH. In fact, in the dsrEFH phylogeny, their genes were positioned in between SAR324 and Nitrospirota (**Figure 4**), indicating a potential horizontal gene transfer of *dsrEFH* from these phyla.

### Members of the transitionary clade encode dsrR, a protein involved in oxidative type dsr

The *dsr* gene content in SAR324 genomes (**Figure 2**) revealed that many of the dsrD*-*encoding SAR324 all also encode dsrR, a protein that has been shown to participate in regulation of sulfur oxidation and correlated with dsrEFH and dsrL expression (33). This observation is intriguing because a combination of oxidative and reductive genes consisting of *dsrD, dsrEFH*, and *dsrR*, has not been described. Therefore, we set out to uncover what other phyla encoded dsrR.

To this end, we searched our dereplicated sulfur cycling database for homologs to dsrR and generated a maximum likelihood dsrR gene tree which was rooted to HesB proteins, which have homologous domains to dsrR (**Figure 6**). Interestingly, besides SAR324 (7 genomes) the only other phyla to encode dsrR were Pseudomonadota (119 genomes), Nitrospirota (3 genomes), and Nitrospinota (1 genome). Nitrospinota was the only other phyla that encoded dsrD, dsrR and dsrE. This observation further emphasizes that these specific members of the transitionary clades likely led to the evolution of oxidative dsr. This data may suggest that these phyla were also involved in the transition to the Pseudomonadota group of oxidative type, given their shared genes, however more data would need to be collected to investigate this claim.

**Figure 6.**
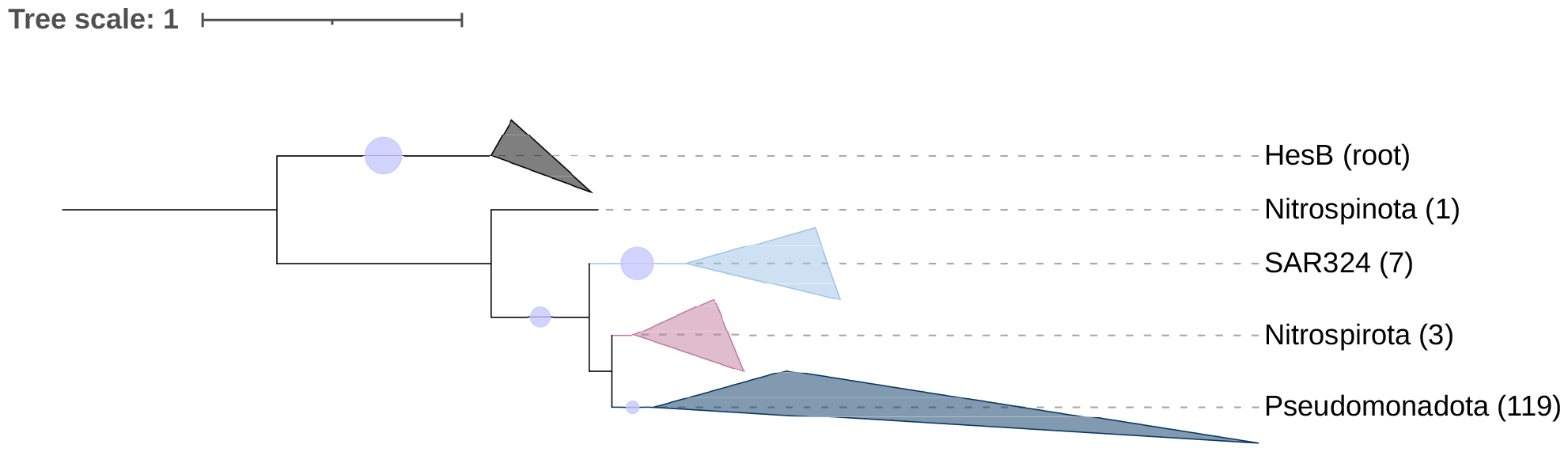
Members of the transitionary clades encode dsrR. Maximum likelihood dsrR tree demonstrates dsrAB encoding phyla also encoding dsrR. The number of genomes is shown in parentheses next to collapsed groups. Bootstrap support >90 is shown with a lavender circle. The tree is rooted to HesB, which has homologous domains to dsrR.

## DISCUSSION

Conducting comparative genomic studies is essential for pinpointing key contributors to global biogeochemical cycles. As the interest in harnessing sulfur-metabolizing bacteria for industrial and bioremediation applications continues to grow, it becomes crucial to identify versatile and pivotal players within the sulfur cycle especially (36,38,70). Further, given that a significant portion of bacteria remain uncultured (71), there is a pressing need for large-scale metagenomics-based investigations to uncover essential targets for cultivation and laboratory experimentation. This study successfully achieves this objective by shedding light on the uncultured candidate phylum SAR324 as a markedly significant participant in sulfur cycling. More specifically, we show how the widespread global and phylogenetic distribution of dsr-encoding SAR324 could lead to influential metabolic impacts across diverse ecological niches.

In addition, the conclusions drawn here are significant because, while attention has been paid to the evolution of the reductive dsr (21,28,72), less is known about the transition to the oxidative type. Through a more in-depth examination of SAR324’s *dsr* gene content and phylogenetic position, we have identified SAR324 as a potential key player in the evolution of novel sulfur metabolism, specifically in the transition to oxidative sulfur metabolism. Additionally, by demonstrating that patterns seen in SAR324 (i.e their unique phylogenetic position in a dsrAB tree, having diverse genomes spanning the oxidative and reductive types, and select genomes carrying genes for both oxidative and reductive dsr) we have shown that members from closely related phyla, such as Nitrospirota, Nitrospinota, Actinomycetota, and CG2-30-57-63, also are particularly important regarding the evolution of the dsr pathway.

Notably, genomes from the phyla Nitrospinota and Actinomycetota stood out as they encoded either colocalized dsrAB, dsrEFH and dsrD or two copies of dsrAB with each being encoded adjacent to dsrD or dsrEFH (**Supplemental Figure 1**). This trait, along with their basal position in the dsrEFH tree (**Figure 4**), raises more evidence that these two phyla are promising candidates for bacteria that were involved in transitioning from the reductive to oxidative type. If the co-expression of dsrEFH and dsrABC is essential for sulfur oxidation, their proximity in the genome may enhance the likelihood that these organisms can perform sulfur oxidation. It is plausible that dsrEFH and a functional oxidative metabolism had to evolve before these genomes could dispense with dsrD and lose the reductive metabolism. We have posited this as a reason these genomes carry genes for both dsrD and dsrEFH. A proposed mechanism for this evolution, as well as a summary of which phyla we have identified that are associated with various *dsr* gene combinations, is illustrated in **Figure 7**.

**Figure 7.**
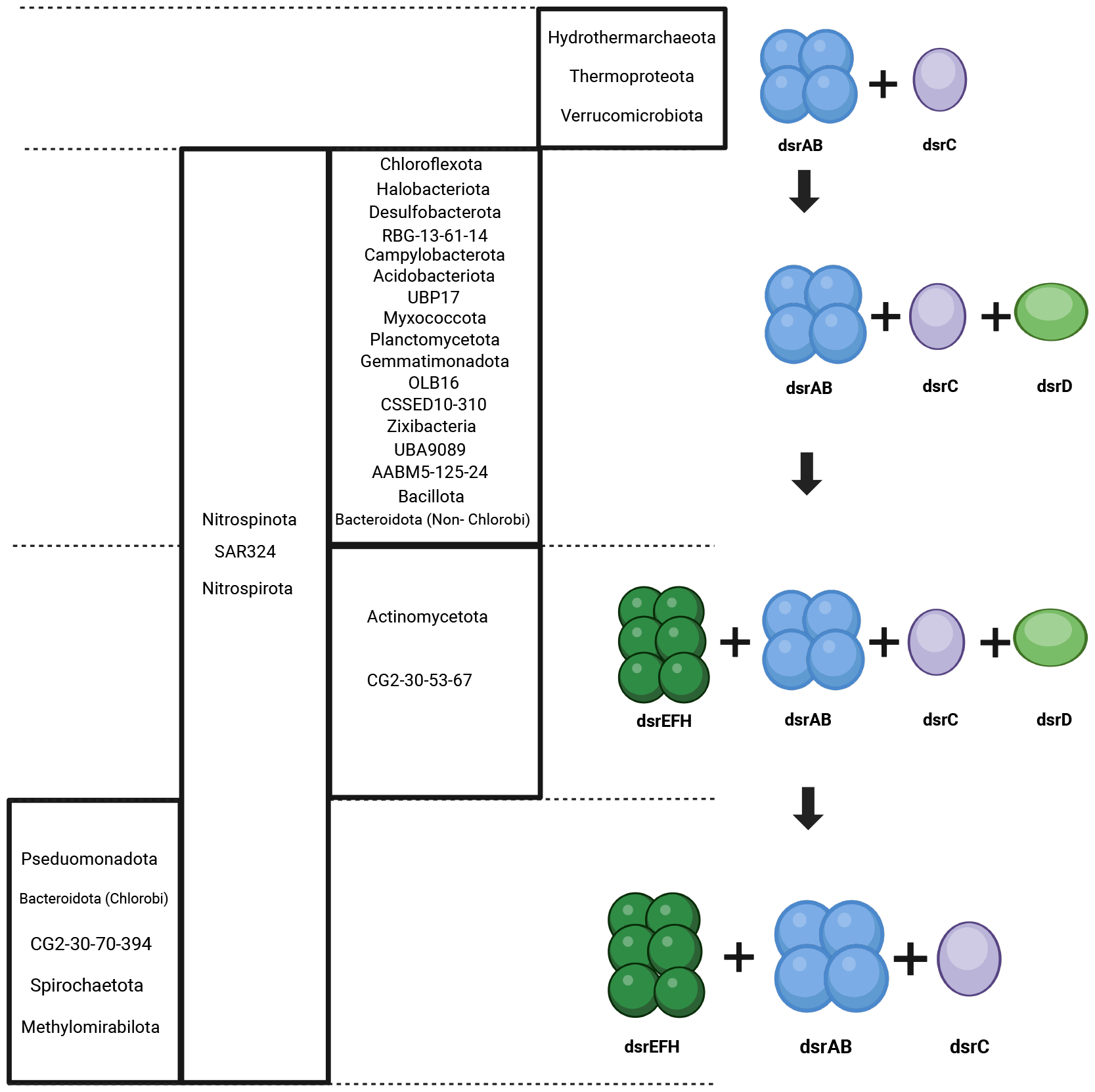
Proposed mechanism for acquisition/loss of key *dsr* genes in the transition from the reductive to oxidative type. Phyla which have genome representatives with the various *dsr* gene combinations are annotated on the illustration. Figure generated with Biorender.com.

In addition to narrowing down the phyla and area of the phylogenetic tree associated with the evolution of oxidative dsr, these observations have implications for future genomics studies, as often studies use solely taxonomic classification to determine what metabolisms are being carried out by certain organisms (73,74). The data presented here clearly reveal that the picture is more complex. This study demonstrates that more thorough metabolic analyses are necessary to fully understand biogeochemical processes that are carried out in by various groups of microorganisms.

Considering these findings, unanswered questions persist, necessitating follow-up studies on microorganisms with genomes placing them in the transitionary clades to unravel their metabolic potential. Transcriptomics and proteomics, coupled with metagenomics, are proposed for future investigations, aiming to determine not only the presence of genes but specifically their transcription. This approach could unveil whether these bacteria activate both pathways simultaneously, or if the presence of certain genes is vestigial. Addressing these questions will contribute to a deeper understanding of the selective pressures governing these genes. Additionally, for phyla like CG2-30-67 and Actinomycetota, where only a few genomes with dsr were found, it underscores the ongoing need for continued sequencing of microbial communities from diverse environments.

Finally, as an ancient metabolism, sulfur cycling, including dsr metabolism, is believed to have played a role in regulating the redox state of early Earth (8). During and post the Great Oxidation Event, microbial metabolism underwent evolution to adapt to the changing environment (75). The dsr pathway, with its ability to reverse direction, may have been crucial is sustaining life and producing energy during this time. Research indicates that certain phyla in the transitionary clades associated with this pathway exhibit remarkable metabolic diversity, enabling them to thrive in various environments (45,76). This feature may have given them the capacity to adapt and shift in response to evolutionary changes, making them intriguing subjects for further investigation from an evolutionary perspective. Unraveling the mechanisms behind this metabolic switch in these phyla becomes particularly intriguing, especially in the context of our own changing climate.

## Data availability

NCBI GenBank and JGI accession numbers for all genomes included in this paper are listed in **Supplemental Tables 1, 2 and 3**. NCBI BioProject numbers for genomes in which metadata was analyzed are listed in **Supplemental Table 1**. The scripts used to visualize gene organization is available at https://github.com/cody-mar10/operon_finder.

## Acknowledgments

We thank the original authors that generated and distributed the publicly available data analyzed in this study. This research was supported by the National Science Foundation under grant numbers DBI2047598 and OCE2049478. CM was funded by a National Science Foundation Graduate Research Fellowship.

## Author contributions

Conceptualization, K.M.K. and K.A.; Methodology, K.M.K., K.A., M.V.L., C.M.; Formal Analysis, K.M.K.; Investigation, K.M.K., M.V.L., and K.A.; Resources, K.A.; Data curation, K.K. and K.A.; Writing — Original Draft, K.K.; Writing — Review & Editing, K.M.K., M.V.L., C.M., and K.A.; Visualization, K.K.; Supervision, K.A.; Project Administration, K.A.; Funding Acquisition, K.A.

## Notes

### Competing Interest Statement

The authors have declared no competing interest.

### Summary of Updates

Updated figures to address manuscript conversion issue; Figure 4 and supplemental figure 1 revised

