## Supplemental Figure 1 for "SAR324 and related lineages are associated with the evolutionary history and origins of dsr-mediated sulfur oxidation"

**AFFILIATIONS:**
[1] Department of Bacteriology, University of Wisconsin-Madison, Madison, WI, 53706, USA
[2] Freshwater and Marine Sciences Program, University of Wisconsin-Madison, Madison, WI, 53706, USA
[3] Microbiology Doctoral Training Program, University of Wisconsin-Madison, Madison, WI, 53706, USA
[4]Department of Integrative Biology, University of Wisconsin-Madison, Madison, WI, 53706, USA

*Correspondence and requests for materials should be sent to K.A

This PDF file includes:

**Supplemental Figure 1. Dsr gene content and arrangement for the transitionary clades**

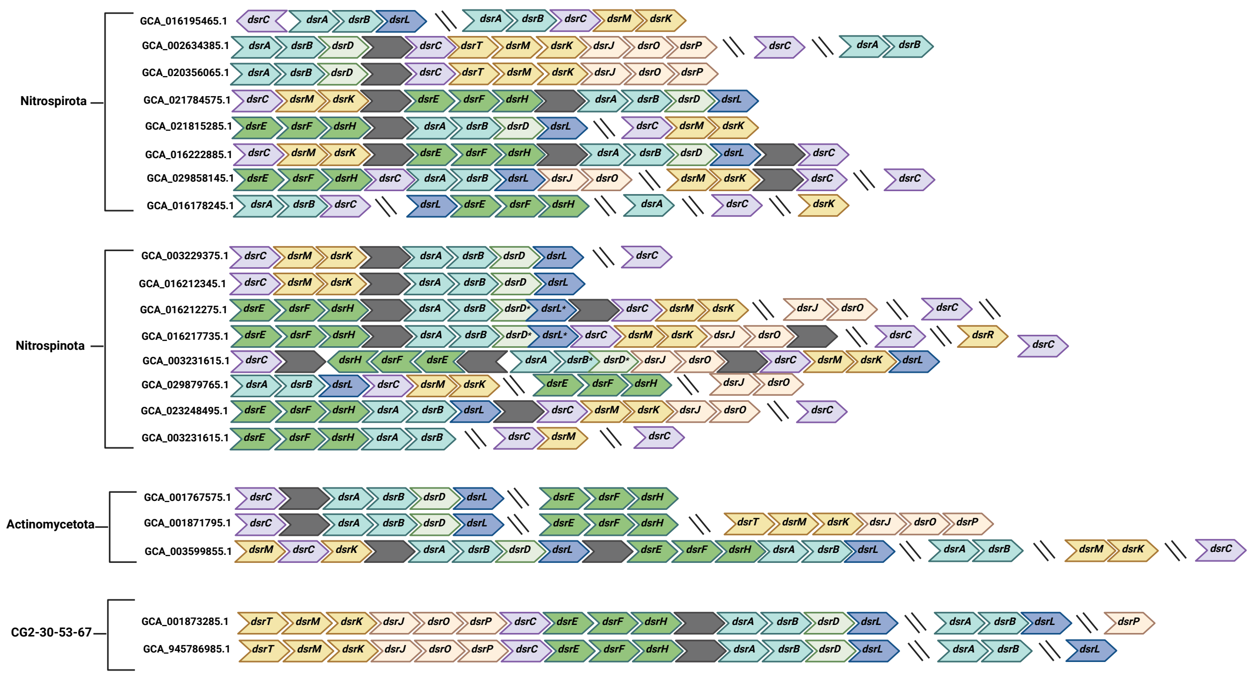


* Proteins found encoded for on same open reading frame

**Supplemental Figure 1. Dsr gene content and arrangement for the transitionary clades.** All dsr-encoding genomes are shown for phyla Actinomycetota and CG2-30-53-67 while representative genomes were chosen for Nitrospinota and Nitrospirota. Dashed lines indicate the end of a scaffold while grey boxes indicate one or more hypothetical or non-dsr proteins. Figure created with BioRender.com
